# Lmo7a Coordinates Neural Crest Migration and Lineage Specification by Regulating Cell Adhesion Dynamics

**DOI:** 10.1101/2020.05.30.125468

**Authors:** David Tatarakis, Adam Tuttle, Thomas F. Schilling

**Author notes:** Author for Correspondence: Thomas F. Schilling, 4109 Natural Sciences II, Department of Developmental and Cell Biology, University of California, Irvine, CA 92617.

## Abstract

Cell migration requires dynamic regulation of cell-cell signaling and cell adhesion. Neural crest (NC) cells are highly migratory cells, which undergo an epithelial-mesenchymal transition (EMT) to leave the neural epithelium and migrate throughout the body to give rise to many different derivatives. We have identified a Lim-domain only (Lmo) protein, Lmo7a, expressed in early NC cells that controls both actin cytoskeletal dynamics and Wnt signaling during NC migration. In embryos deficient in Lmo7a, many NC cells fail to migrate away from the dorsal midline, and form aggregates. Unlike the majority of NC cells that appear to migrate normally, cells that aggregate in Lmo7a-deficient embryos mislocalize paxillin (Pxn) and have reduced levels of phosphorylated focal adhesion kinase (pFAK). Lmo7a loss-of-function also disrupts canonical Wnt signaling such that after the onset of NC cell migration, Wnt responses and nuclear β-catenin levels increase in the cells that aggregate. However, this increase in Wnt signaling appears secondary to the defect in migration. Similar to mutants for other Wnt regulators in NC cells, the NC cells in Lmo7a-deficient aggregates exhibit gene expression signatures of pigment cell progenitors, but also express markers of Schwann cell progenitors, suggesting a role for Lmo7a in pigment-glial specification. We propose that Lmo7a modulates cell adhesion to facilitate both robust NC cell migration and a subset of lineage decisions.

## INTRODUCTION

During embryonic development, cell migration and lineage specification must be tightly coordinated. Many of the mechanisms driving progenitor cell migration also regulate differentiation toward specific lineages (McBeath et al, 2004; He et al, 2018). In vertebrates, this is particularly true for mesenchymal cell populations such as migratory neural crest (NC) cells, which emerge from the neural ectoderm and disperse to generate an extraordinary variety of cell types throughout the body including neurons, glia, pigment cells, cartilage, and bone. Despite extensive studies of both intrinsic and extrinsic factors that specify these different fates, how the acquisition of distinct NC lineages relate to their migratory behaviors remains largely unclear (Kalcheim and Kumar, 2017).

NC cells are induced at the neural plate border during neural tube closure through a combination of Wnt, BMP, FGF, and Notch signaling (Stuhlmiller and Garcia-Castro, 2012). They subsequently undergo epithelial-mesenchymal transition (EMT) and migrate away from the dorsal midline along distinct trajectories throughout the embryo (Kerosuo and Bronner-Fraser, 2012; Mayor and Theveneau, 2013). Among these migratory paths, are cranial NC cell streams that populate the pharyngeal arches (PAs) to form the facial skeleton, as well as distinct lateral and medial pathways in the trunk in which NC cells form pigment cells in the skin or sensory neurons and glia in peripheral nerves, respectively. Some in vivo lineage tracing and in vitro clonal analyses have suggested that these fates depend entirely on the migratory environments and final destinations of NC cells (Fraser and Bronner-Fraser, 1991; Dupin et al, 2010; Baggiolini et al, 2015). Other experiments have provided evidence for early lineage specification in premigratory NC and a link between initial position, migratory path, and cell fate (Stemple and Anderson, 1992; Schilling and Kimmel, 1994; Krispin et al 2010). In recent years, the advent of single cell transcriptomics has given us a more detailed picture of the degree to which NC cell fates are both dynamic and heterogeneous during migration (Morrison et al, 2017; Lukoseviciute et al, 2018; Soldatav et al, 2019).

Canonical Wnt signaling plays important roles in inducing NC cells, promoting their migration, and driving lineage decisions at later stages. Tight regulation of signaling levels is vital for proper initiation of migration (Maj et al, 2016; Hutchins and Bronner, 2018; Ahsan et al., 2019) and also biases cells toward pigment versus glial cell fates through regulation of genes such as *Sox10/Foxd3* and *Pax3/7*, respectively (Dorsky et al., 1998; Minchin and Hughes, 2008; Curran et al, 2010). We previously demonstrated novel roles for Ovol1a and Rbc3a/Dmxl2 in promoting NC migration, due at least in part to changes in responses to Wnt signaling (Piloto and Schilling, 2010; Tuttle et al, 2014). Both are specifically expressed in premigratory NC cells, and loss-of-function of either gene disrupts the migration of subsets of NC cells, which form aggregates in the dorsal midline and acquire pigment cell fates. Ovol1a is a direct Wnt target (Li et al, 2002), while Rbc3a/Dmxl2 controls Wnt receptor trafficking, and both alter the localization of cadherins (such as Cdh2) important for migration. These results establish a link between adhesive mechanisms that govern NC migration, Wnt signaling, and the decisions that lead toward specific cell fates.

In a microarray screen to identify genes downregulated in *tfap2a/g*-deficient zebrafish embryos, we discovered the Lim-domain-only 7 gene *lmo7a* (Hoffman et al. 2007, Tuttle et al., 2014). Lim-domain proteins vary in structure and cellular function and include the Lim-domain-only (Lmo) subclass. Lmo1-4 contain no annotated functional domains apart from multiple Lim domains and function as nuclear transcriptional co-regulators important in cancer progression (Matthews et al, 2013; Sang et al, 2014). Lmo4 promotes EMT of NC and neuroblastoma cells through direct binding to Snail and Slug transcription factors (Ochoa et al, 2012; Ferronha et al, 2013). A protein containing four and one-half Lim domains (FHL2) interacts with β-catenin (β-cat) to either increase or decrease levels of TCF/LEF dependent transcription, depending on the cellular context (Martin et al, 2002; Hamidouche et al, 2008). Other Lim-domain family members contain multiple functional domains and have more divergent roles.

Lmo7, despite its name, contains a calponin homology (CH) domain, a PDZ domain, and a single Lim domain. In mammals, multiple functions have been described for Lmo7, including promoting expression of myogenic transcription factors in skeletal muscle cells (Holaska et al, 2006; Dedeic et al, 2011) and regulating afadin-nectin-E-cadherin junctions in epithelial cells (Ooshio et al, 2004) and in the cuticular plate of the cochlea (Du et al, 2019). Interestingly, its function in myogenesis requires entry into the nucleus and transcriptional regulation, while its epithelial functions involve interactions with the actin cytoskeleton and membrane-associated proteins. Lmo7 also influences cancer cell metastasis, such as the expression of myocardin-related transcription factors through regulation of Rho-dependent actin dynamics at the cell membranes of breast cancer cells (Nakamura et al, 2005; Hu et al, 2011, Teixeira et al, 2014). A paralog of Lmo7, LIMCH1, regulates cell migration through roles in focal adhesion (FA) formation and actomyosin dynamics (Lin et al, 2017). Lmo7 can localize to FAs and act as a shuttling protein to mediate integrin signaling in HeLa cells and mouse embryonic fibroblasts (Holaska et al, 2006; Wozniak et al, 2013). Both LIMCH1 and Lmo7 are associated with poor prognosis in human lung cancer (Karlsson et al, 2018), and expression of Lmo7 (also called PCD1) is associated with increased metastasis in numerous human cancers (Kang et al, 2000; Furuya et al, 2002; Sasaki et al, 2003).

Here, we show that zebrafish Lmo7a promotes NC migration and modulates lineage decisions through interactions with canonical Wnt signaling, similar to Ovol1a and Rbc3a/Dmxl2. Lmo7a is expressed in premigratory NC cells where it localizes to cell membranes, and loss-of-function leads to aggregation of subsets of NC cells at the dorsal midline. These cells show elevated nuclear β-cat as well as altered paxillin (Pxn) localization and reduced phosphorylated focal adhesion kinase (pFAK). Furthermore, analysis of gene expression in these NC cell aggregates reveals that the cells adopt identities of pigment and glial progenitors, but not other NC lineages. Our results suggest that Lmo7a has a dual role in promoting migration of NC cells and regulating lineage decisions through modulation of canonical Wnt signaling and cell adhesion dynamics.

## RESULTS

### *Lmo7a* is expressed in and required for migration of subsets of NC cells

Zebrafish *lmo7a* was identified in a microarray screen of *tfap2a/g*-deficient embryos, which lack NC cells (Hoffman et al., 2007; Tuttle et al., 2014). Whole mount in situ hybridization (ISH) first detected *lmo7a* expression in cranial NC cells at 12 hours post-fertilization (hpf), just prior to the onset of NC cell migration (Figure 1A). Expression persisted at 16 hpf in migrating NC cells in the PAs and between the eyes (Figure 1B), but was no longer detected at 24 hpf. At later embryonic stages (48-72 hpf), expression was restricted to the notochord and somite boundaries (Figure 1C, D).

**Figure 1:**
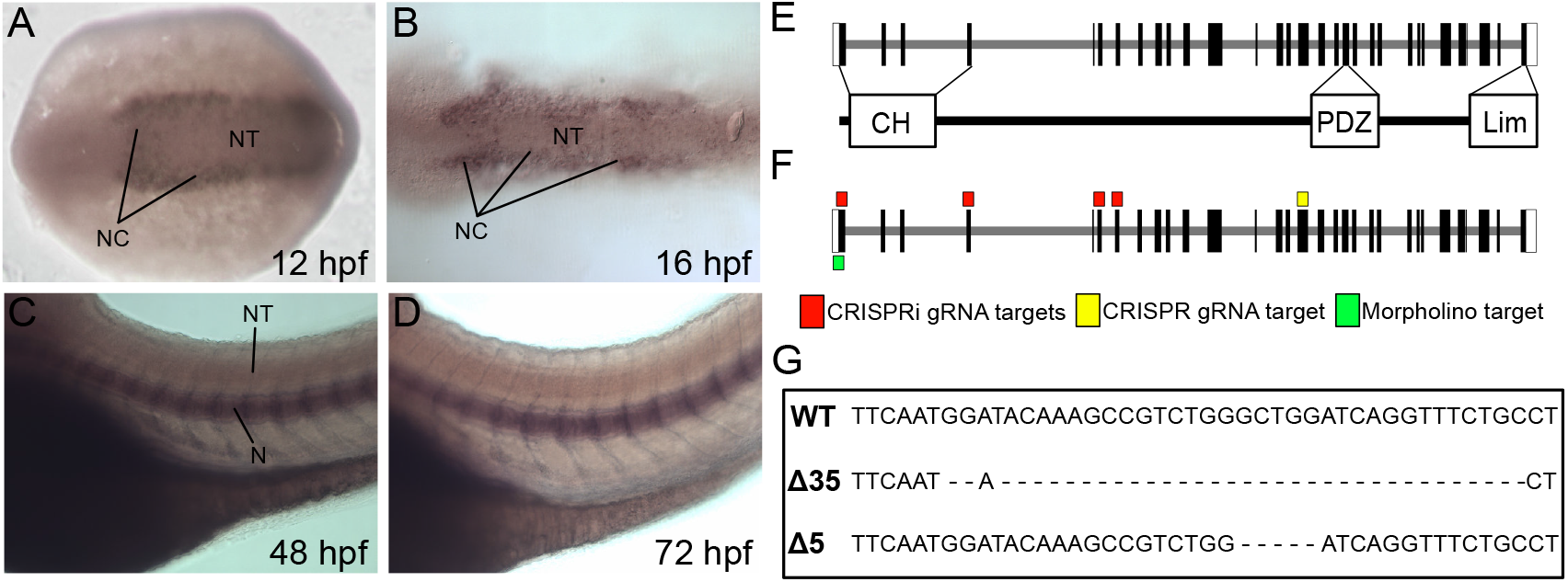
*lmo7a* expression and domain structure. **(A-B)** Whole mount in situ hybridization (ISH) for *lmo7a* mRNA (dorsal views, anterior to the left). Expression in premigratory NC cells at 12 hpf (**A**) and migratory NC cells at 16 hpf (**B**). (**C-D**) Whole mount ISH at 48 hpf (**C**) and 72 hpf (**D**) (lateral views), shows expression in the notochord and somite boundaries. (**E**) The *lmo7a* genomic locus consists of 33 exons. The full-length protein contains calponin homology (CH), PDZ, and Lim domains. (**F**) An antisense morpholino targeted the first exon, while CRISPR and CRISPRi gRNAs targeted exons 1,4,6, and 7. (**G**) Sequences for two CRISPR deletion alleles generated in Exon 16 compared to WT sequence.

Lmo7a contains CH, PDZ and Lim domains (Figure 1E). Two deletions in *lmo7a*, −5 bp and −35 bp, were generated by CRISPR-Cas9 gene editing using a guide RNA targeting exon 16 just upstream of the PDZ domain (Figure 1F, G). Trans-heterozygous mutants carrying the *Tg(-4.9sox10:nEOS)* transgene (hereafter referred to as sox10:nEOS, which labels the nuclei of pre-migratory and migrating NC cells) appeared largely normal but NC cells formed small aggregates (5-10 cells/aggregate; ~10-20 cells/embryo) at the dorsal midline of the neural tube extending along the anterior-posterior (A-P) axis from the midbrain-hindbrain boundary to the anterior spinal cord (Figure 2A,B). Such aggregates closely resemble the phenotypes of *rbc3a*^-/-^ and *ovol1a*-deficient embryos (Piloto et al, 2010; Tuttle et al., 2014).

**Figure 2:**
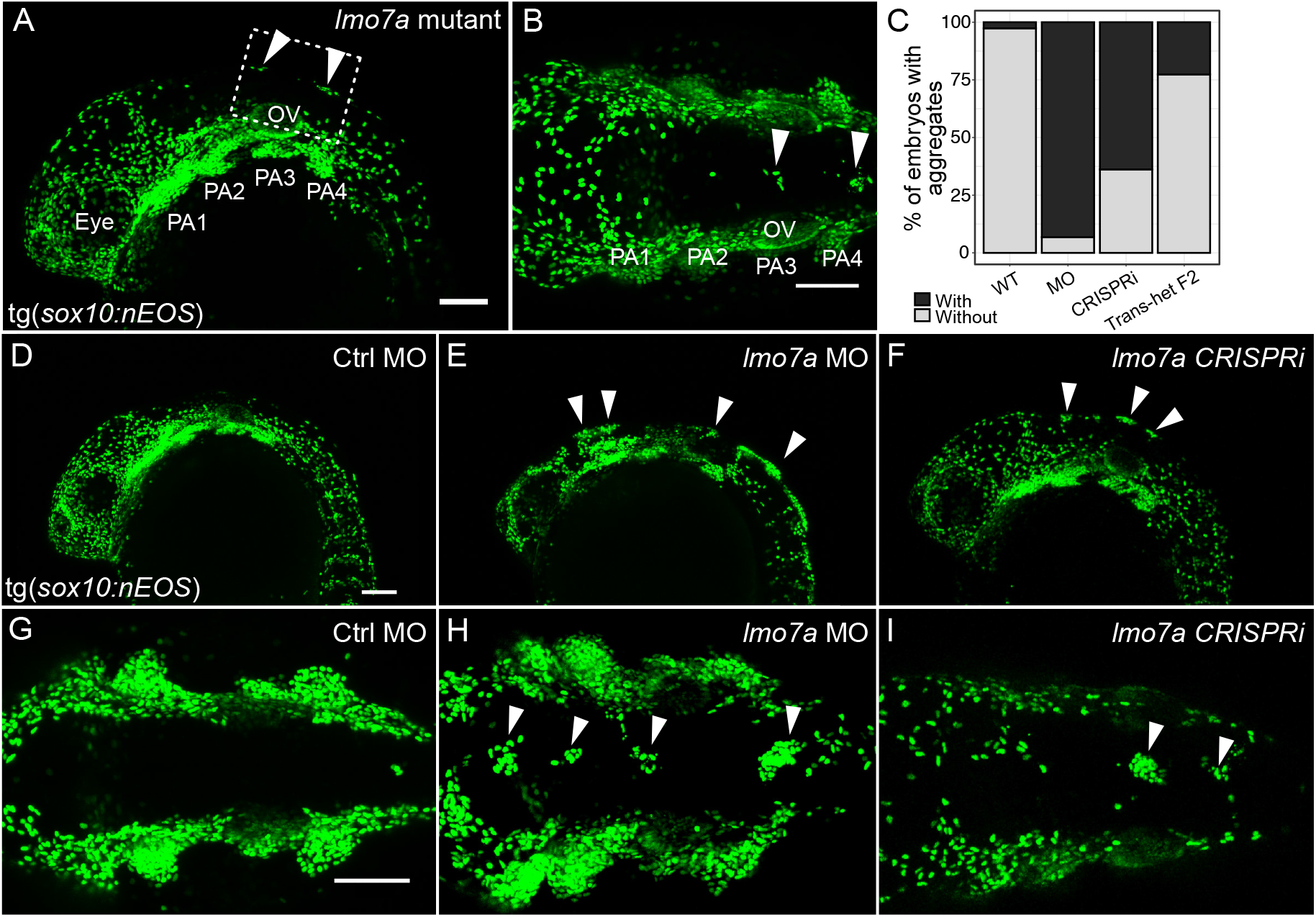
*lmo7a* knockdown disrupts migration of subsets of NC cells. **(A-B, D-I)** Whole mount live confocal images of 24 hpf *sox10:nEOS* embryos. **(A-B)** NC aggregates form along the dorsal midline in *lmo7a* trans-heterozygous mutants, in contrast to WT siblings. **(C)** Percentages of embryos displaying >10 NC cells at the dorsal midline in various *lmo7a* gene perturbations. **(D,G)** Embryos injected with 4 ng of control morpholino (MO), **(E,H)** Embryos injected with 4 ng of antisense MO targeting *lmo7a* **(F,I)** Embryos injected with *dCas9* mRNA and 4 gRNAs targeting the coding region of *lmo7a*. NC cell aggregates at the dorsal midline (arrowheads). **(A, D-F)** Lateral views. **(B, G-I)** Dorsal views. Scale bars = 100 μm **(A,B,D,G)**. PA=Pharyngeal Arch, OV=Otic Vesicle

Similar to *lmo7a* mutants, knockdown of *lmo7a* using an antisense morpholino targeting the translation start site (*lmo7a*-MO) (Figure 1H) in sox10:nEOS fish resulted in NC aggregates (5-30 cells/aggregate; ~50-100 cells/embryo) at the dorsal midline in ~93% of injected embryos compared to sibling embryos injected with a control MO (Figure 2D-E,G-H). Aggregates became distinct by 18 hpf while other surrounding NC cells migrated away from them and ventrally into the PAs at approximately the same rate as WT cells (Suppl Movie 1-2). CRISPR inhibition (CRISPRi) produced similar NC cell aggregates in ~64% of injected embryos (Figure 2F,I). With CRISPRi, expression of *lmo7a* at 12 hpf was nearly undetectable (Figure S1). These results provide independent confirmation that Lmo7a function is required for subsets of NC cells to migrate.

### Lmo7a localizes to NC cell membranes and its function requires the calponin homology domain

Lmo7 was previously shown to function at the membrane in epithelial cells where it interacts with adherens junctions and/or focal adhesions (FAs), and in the nucleus in muscle cells, where it binds the transcription factor Emerin (Ooshio et al, 2004; Wozniak et al, 2013; Holaska et al, 2006). To determine the subcellular localization of Lmo7a in NC cells, we generated a fusion construct encoding superfolder GFP (sfGFP) fused to the N-terminus of Lmo7a, *sfGFP-lmo7a*. This mRNA was injected at the 1-cell stage into *Tg(-4.9sox10:lyn-tdTomato*) embryos (hereafter referred to as *sox10:lyn-tdTom*) to mark NC cell membranes. At 18 hpf, the *sfGFP-lmo7a* fusion protein was restricted to bright puncta that co-localized with *sox10:lyn-tdTom*, with little to no expression detected in cell nuclei (Figure 3A-C), indicating a potential role at the membrane.

**Figure 3:**
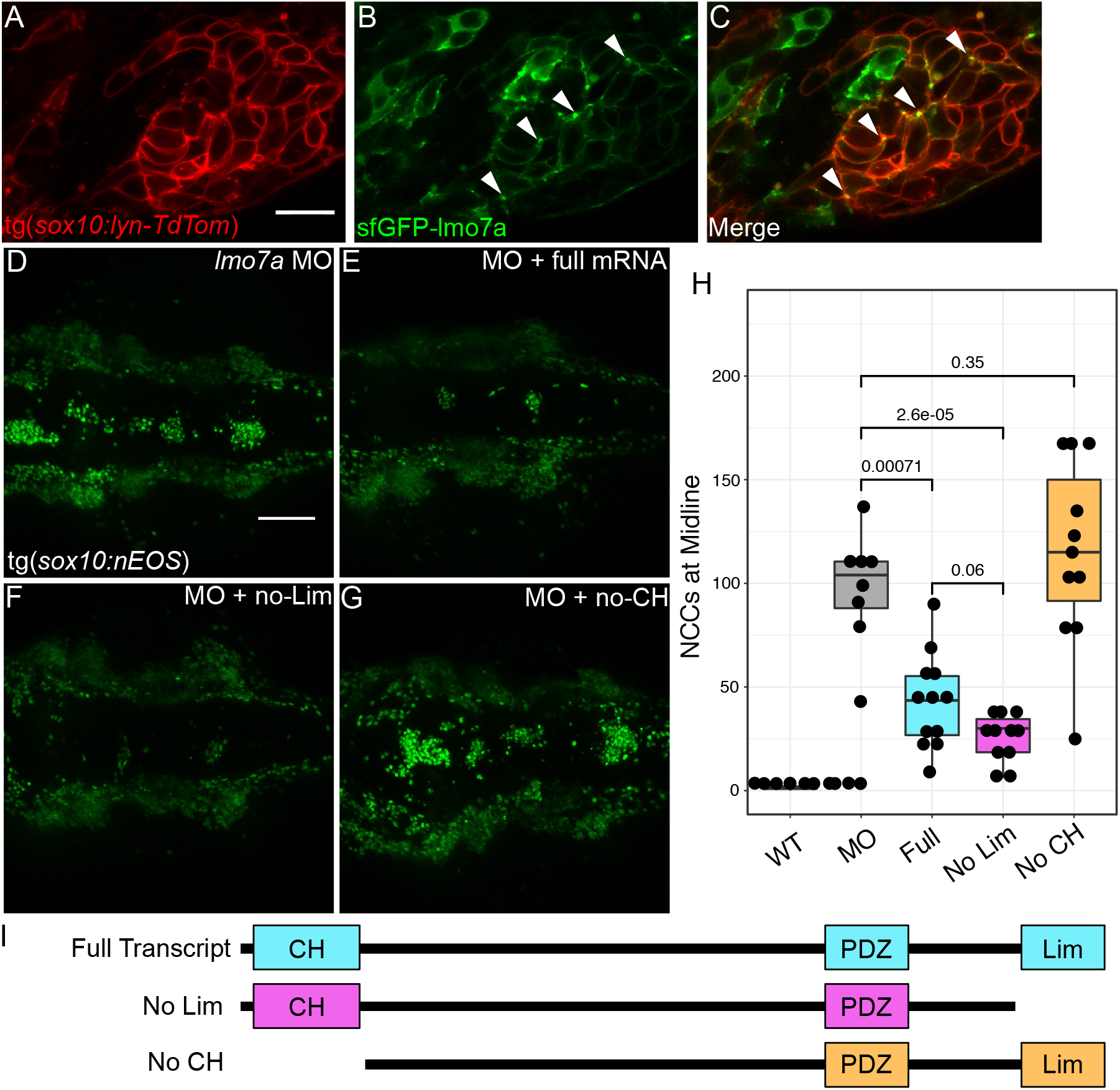
Lmo7a localizes to the plasma membrane and requires the CH domain in NC. **(A-C)** Whole mount live confocal images of WT *sox10:lyn-TdTomato* embryos injected with *sfGFP-lmo7a* fusion mRNA. sfGFP-Lmo7a puncta at the plasma membrane of NC cells (arrowheads).(lateral views, anterior to the left). **(D-G)** Whole mount live confocal images of 24 hpf *sox10:nEOS* embryos (dorsal views, anterior to the left). **(D)** *lmo7a* MO-injected embryos display large NC aggregates. Both full length mRNA **(E)** and mRNA lacking the Lim domain **(F)** reduce the number of NC cells in aggregates, while mRNA lacking the CH domain **(G)** does not. **(H)** Quantification of phenotype severity based on median number of NC cells at midline/embryo (n = 8 embryos per condition). **(I)** Schematics of rescue transcripts. *Tg(sox10:lyn-tdTomato)* For **(H)**: Medians: MO = 104 cells/embryo, MO+full mRNA = 43.5 cells/embryo, MO+noLim mRNA = 30 cells/embryo, MO+noCH mRNA = 115 cells/embryo. Line indicates median, boxes indicate IQR, whiskers indicate IQR*1.5, point indicates outlier. Kruskal-Wallis ANOVA: p-value <0.0001. Posthoc Wilcoxon tests (BH corrected) p-values: ***<0.001; ****<0.0001, ns>0.05. Scale bars = 20 μm **(A)** and 100 μm **(D)**.

Lmo7a contains CH, PDZ and Lim domains, all implicated in protein-protein interactions (Figure 1G). To test requirements for these domains in NC cell migration, rescue experiments were performed in an *lmo7a*-MO background. Co-injection of *lmo7a*-MO with mRNA encoding full-length *lmo7a* significantly reduced the severity of the migration phenotype as quantified by the number of NC cells that failed to migrate away from the midline by 24 hpf (MO median = 104 cells/embryo, MO+*lmo7a*-full median = 43.5 cells/embryo; p<0.001) (Figure 3D, E, H). Next, mRNAs encoding full-length *lmo7a* lacking either the Lim or the CH domain were co-injected with *lmo7a*-MO. Interestingly, removal of the Lim domain caused no significant change in the ability of injected *lmo7a* mRNA to rescue the *lmo7a*-MO phenotype (MO+*lmo7a*-*noLim* median = 30 cells/embryo; p<0.0001) while *lmo7a* lacking the CH domain failed to rescue (MO+*lmo7a*-full median = 115 cells/embryo; p=0.35) (Figure 3F, G, H). The CH domain mediates interactions with the actin cytoskeleton. These data suggest that the CH domain but not the Lim domain is required for *lmo7a*’s function in early NC cell migration, further supporting a role for Lmo7a at the membrane in NC cells and potential interactions with the cytoskeleton.

### NC cells in *lmo7a*-deficient embryos have aberrant accumulation of focal adhesion components

Based on its membrane localization in NC cells and known roles in FAs (Holaska et al, 2006; Wozniak et al, 2013), we hypothesized that Lmo7a likely plays a role in FAs essential for proper filopodial dynamics and migration. To test this idea we next examined filopodial extension and FA formation. We utilized a double transgenic line expressing both a fluorescent paxillin (Pxn) fusion protein under the control of a ubiquitous promoter, *Tg(β-actin:Pxn-EGFP)* as well as *sox10:lynTdTomato* to facilitate live, time-lapse imaging of Pxn-based adhesion complexes in NC cells. Embryos were imaged from 16-19 hpf, when cranial NC cells migrate into the PAs. In WT embryos, NC cells dynamically change morphology as they migrate, extending and retracting filopodia. Transient accumulations of Pxn-EGFP were visible along the membranes of these NC cells during migration (Figure 4A-C). In contrast, in *lmo7a*-deficient embryos, NC cells in aggregates that remained at the midline maintained a rounded morphology with few to no filopodial projections and contained large aggregates of Pxn-EGFP that appeared cytoplasmic (Figure 4D-F; Suppl Movie 3). To investigate if Lmo7a colocalizes with Pxn, we injected *mCherry-Lmo7a* mRNA into WT *tg(B-actin:Pxn-EGFP)* embryos. Overlap of mCherry-Lmo7a puncta and Pxn-EGFP puncta was observed at the membranes of migratory WT NC cells (Figure 4G-I). These observations suggest that in *lmo7a*-deficient embryos, Pxn-FA complexes fail to form properly at the membrane and that NC cells consequently fail to extend directional projections that facilitate migration.

**Figure 4:**
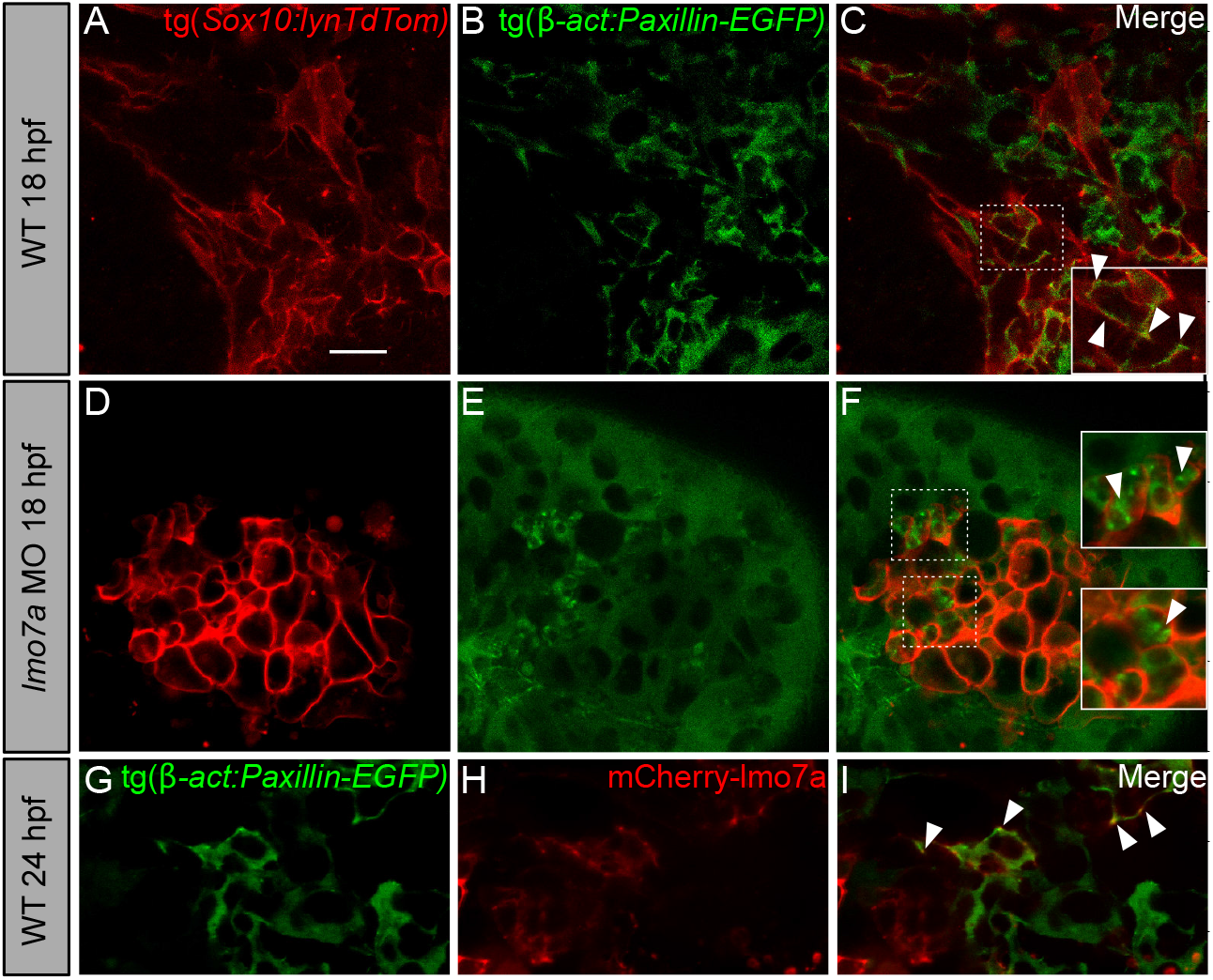
*lmo7a* knockdown results in abnormal NC cell morphology and aggregation of paxillin complexes. **(A-F)** Whole mount live confocal images of *Tg(sox10:lyn-tdTomato; β-actin:paxillin-EGFP)* double transgenic embryos. **(A-C)** NC cells at 18 hpf display long cytoplasmic protrusions (filopodia) and localization of Pxn-EGFP along the plasma membrane (arrowheads) in WT. **(D-F)** NC cells in dorsal midline aggregates in embryos injected with *lmo7a*-MO. Cells are rounded and accumulate Pxn-EGFP in the cytoplasm. **(G-I)** Whole mount live confocal images of transgenic *β-actin:paxillin-EGFP* embryos injected with *mCherry-lmo7a* mRNA. mCherry-Lmo7a puncta colocalize with Paxillin-EGFP in WT NC cells at 24 hpf **(I)** (arrowheads). Scale bar = 15 μm

To investigate if this Pxn accumulation is indicative of changes in FA dynamics, we examined localization of phosphorylated Focal Adhesion Kinase (pFAK) using a polyclonal anti-pFAK (pY576) antibody. At 18 hpf, many bright puncta can be seen along the membranes of migrating NC cells in embryos injected with a control MO (Ctrl MO) (Figure 5A-D). While some puncta can be seen in the membranes of midline aggregate NCs in *lmo7a*-MO-injected embryos (Figure 5E-H), the number of puncta per cell was significantly lower than controls (Ctrl MO mean = 13.1/cell/embryo, *lmo7a* MO mean = 6.92/cell/embryo, p=0.0029) (Figure 5I).

**Figure 5:**
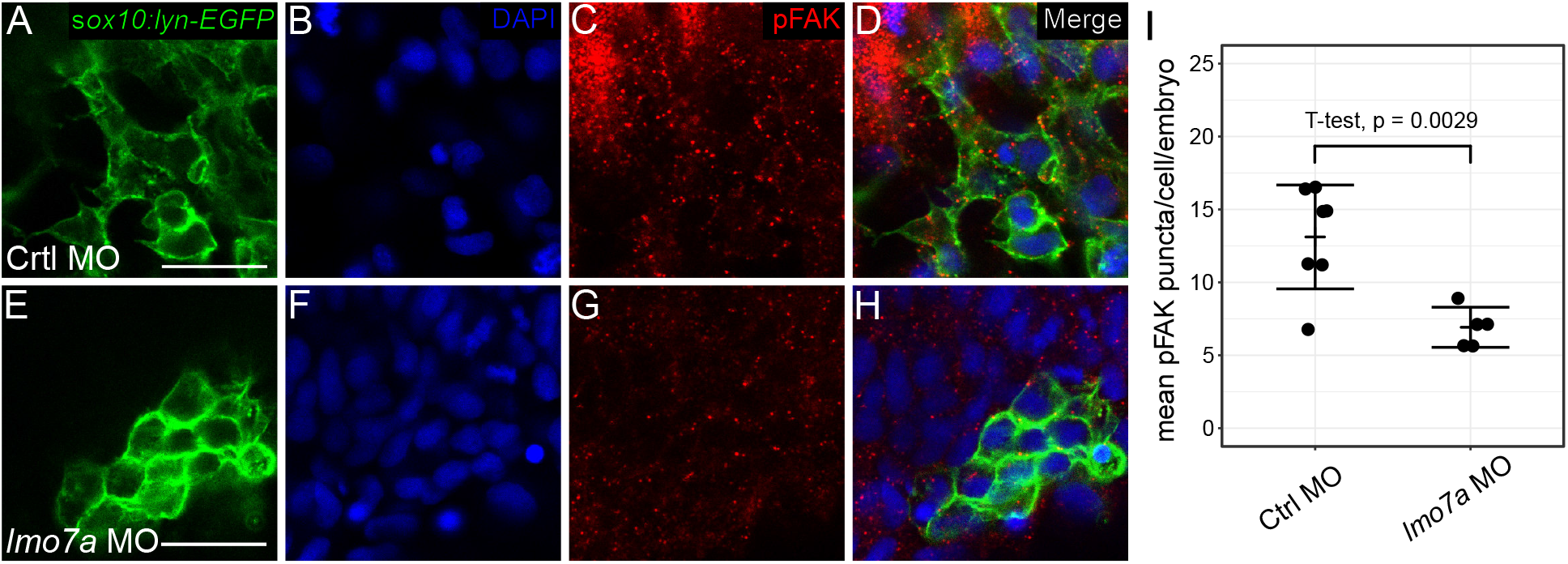
*lmo7a*-deficient NC aggregates have decreased levels of phosphorylated focal adhesion kinase. **(A-H)** Whole mount confocal images of immunostaining with anti-pFAK antibody (red) and DAPI (blue) in *sox10:lynEGFP* transgenic embryos (green). **(A-D)** Migratory cells in Ctrl MO-injected embryos show many bright pFAK puncta along their membranes. **(E-H)** Aggregate NC cells in *lmo7a* MO-injected embryos show fewer pFAK puncta in their membranes. (I) Dots represent the mean number of puncta per cell in 5 cells per embryo. (Ctrl MO n=7 embryos, mean=13.1 puncta/cell/embryo; *lmo7a* MO n=5 embryos, mean=6.2 puncta/cell/embryo). T-test p-value = 0.0029. Line indicates mean. Error bars indicate ±SD. Scale bars = 20 μm.

### Lmo7a deficiency elevates canonical Wnt signaling in NC cell aggregates

We have previously shown that Ovol1a and Rbc3a/Dmxl2 regulate NC cell migration, at least in part, by regulating canonical Wnt signaling (Piloto et al. 2010; Tuttle et al. 2014). To investigate if the NC migratory defects observed in *lmo7a*-deficient embryos also alter Wnt signaling, we examined a canonical Wnt reporter line, *Tg(7xTCF:EGFP)*. In WT embryos, cranial NC cells in PA1 and PA2 as well as over the midbrain expressed the Wnt reporter at 24 hpf (Figure 6A-C). In contrast, most of the NC cells within aggregates in *lmo7a*-deficient embryos were TCF-GFP-positive, regardless of their A-P position (Figure 6D-F). Levels of Wnt responses varied dramatically between cells within an aggregate (Figure 6G-I).

**Figure 6:**
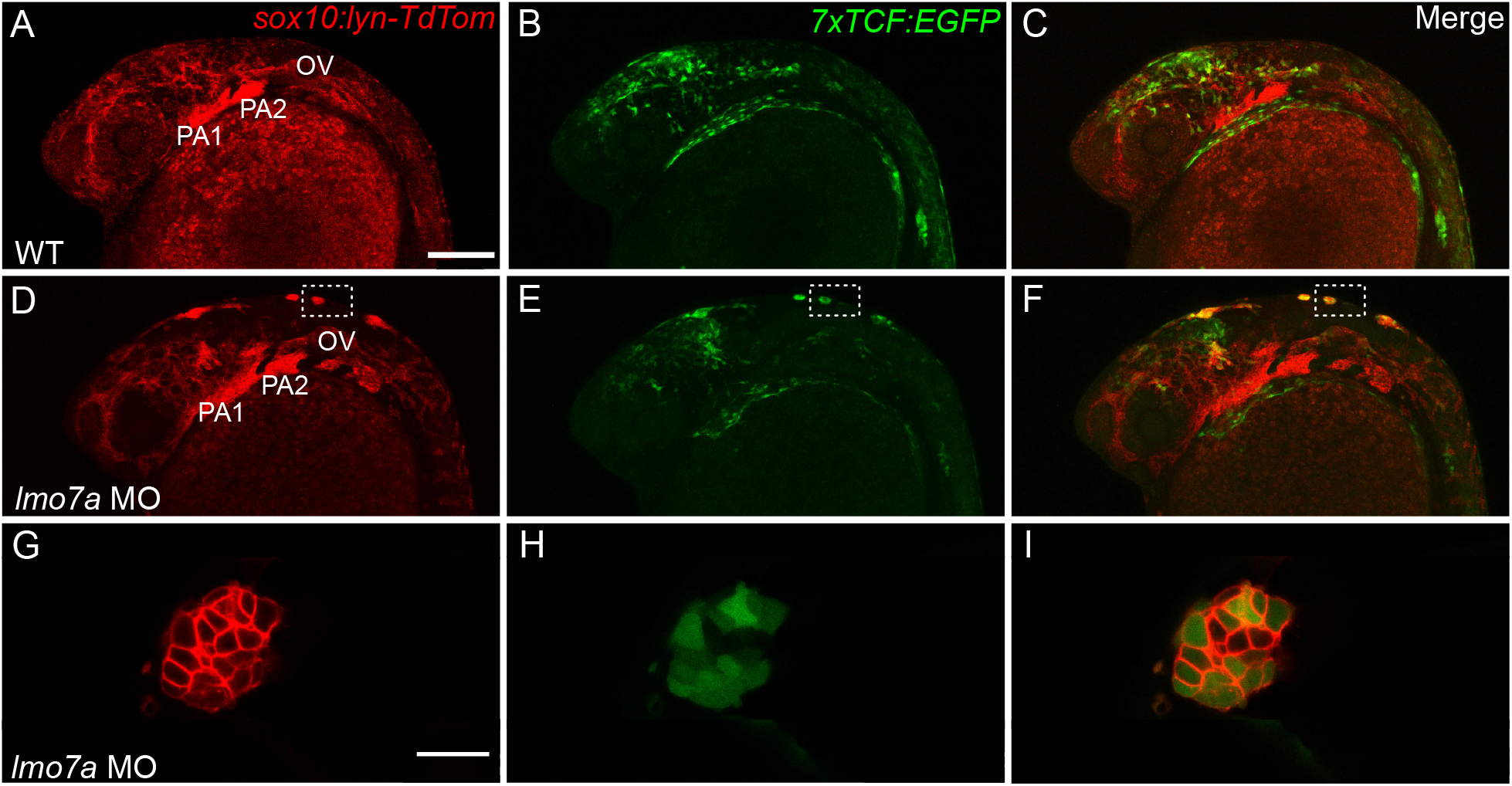
*lmo7a* knockdown increases canonical Wnt signaling in NC midline aggregates. Whole mount live confocal images of double transgenic *Tg(sox10:lyn-tdTomato; 7XTCF:EGFP)* embryos at 24 hpf. **(A-C)** In WT embryos, EGFP is detected in migratory NC cells around the midbrain, anterior hindbrain and PAs 1 and 2. **(D-F)** In *lmo7a* MO injected embryos EGFP is detected in NC aggregates at the dorsal midline regardless of anterior-posterior position. **(G-I)** At higher magnification, most cells within each aggregate are positive for EGFP. **(A-F)** Lateral views. **(G-I)** Dorsal view. Scale bars = 150 μm **(A-C)** 25 μm **(G-I)** and 15 μm **(J-N)**. PA=Pharyngeal Arch, OV=Otic Vesicle

To confirm the apparent increase in canonical Wnt signaling, we analyzed the subcellular localization of β-cat in WT and *lmo7a*-deficient NC cells at 24 hpf (Figure S2A-H). Similar to our previous results for Ovol1a and Rbc3a, in *lmo7a*-deficient embryos, NC cells in the midline aggregates displayed increased levels of β-cat in the nucleus as compared to WT NC cells (Figure S2I).

### *lmo7a*-deficient NC cell aggregates co-express pigment and glia markers

Increased canonical Wnt signaling can drive NC cells toward pigment cell fates (Curran et al, 2010). To determine if this is the case in NC aggregates in *lmo7a*-deficient embryos, we performed in situ Hybridization Chain Reaction (HCR) for genes that mark different NC lineages at 24 hpf (Figure 7). In *lmo7a*-MO injected embryos, expression of the skeletogenic marker *dlx2a* was restricted to the PAs, similar to WT, and was excluded from midline NC aggregates (Figure 7B, F). In contrast, all of these aggregates expressed *mitfa*, which labels melanocyte progenitors (Figure 7C, G). These results suggest that elevated canonical Wnt signaling in NC cells that aggregate at the midline in *lmo7a*-deficient embryos promotes their differentiation as pigment cells, similar to our previous results for Ovol1a and Rbc3a/ Dmxl2.

**Figure 7:**
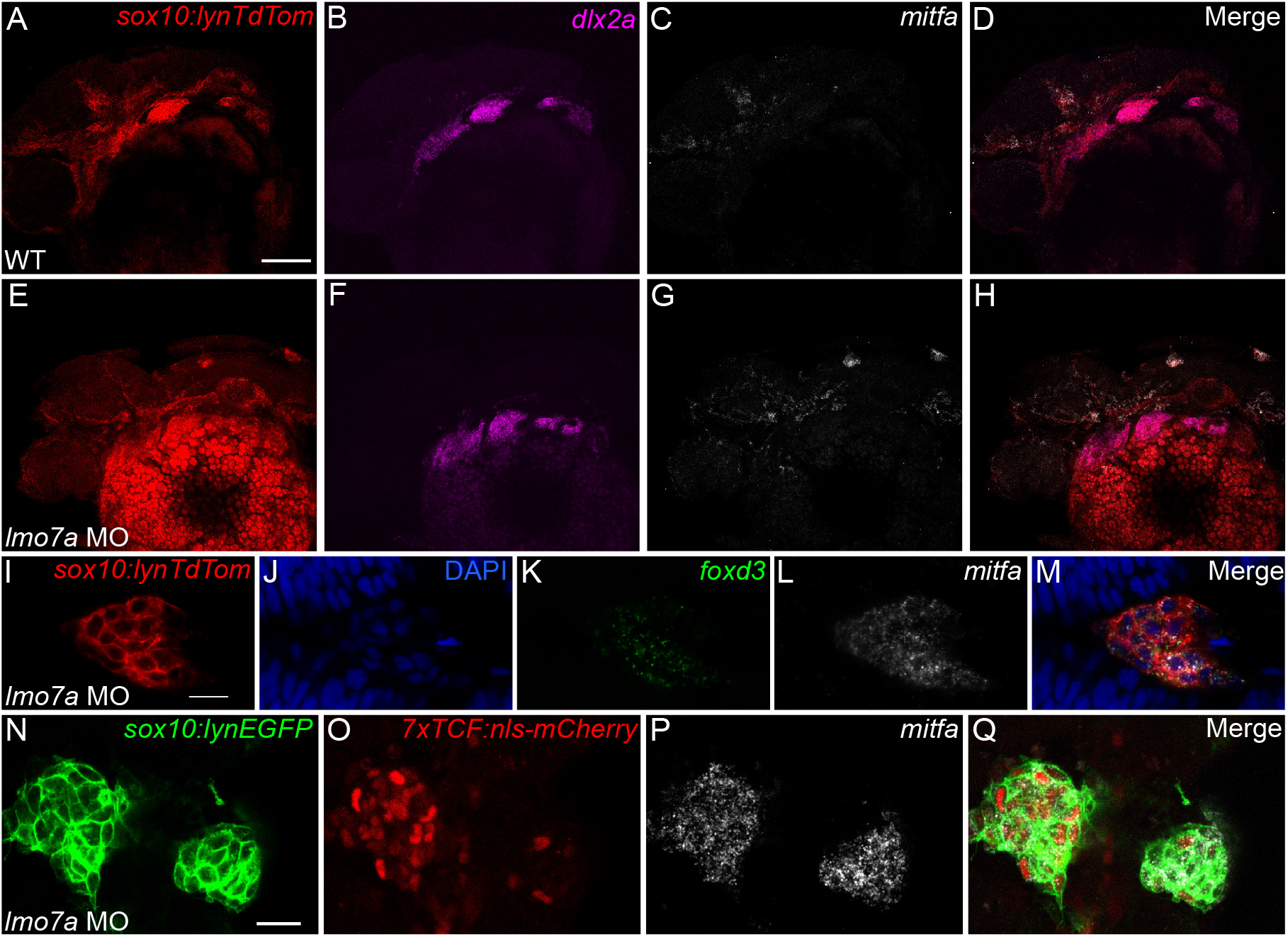
NC aggregates adopt a bipotential pigment/glial cell fate. **(A-H)** Whole mount confocal images of in situ hybridization chain reaction (HCR) for NC lineage markers in transgenic *Tg(sox10:lyn-tdTomato)* embryos. **(A-D)** In WT embryos, *dlx2a* is expressed in NC cells within the PAs and *mitfa* expression is observed in many NC cells outside the PAs. **(E-H)** In *lmo7a* MO-injected embryos, *dlx2a* expression is unaffected, but *mitfa* expression marks NC aggregates at the dorsal midline. **(I-M)** Whole mount confocal images of HCR for NC lineage markers. **(I)** NC aggregates in *lmo7a* MO-injected embryos express both the glial progenitor marker *foxd3* **(J)** and the melanocyte progenitor marker *mitfa* **(L)**, and some cells co-express both markers (M). **(N-Q)** Whole mount confocal images of HCR for NC lineage markers in *lmo7a* MO-injected double transgenic *Tg(sox10:lyn-EGFP; 7XTCF:nls-mCherry)*. Both mCherry+ and mCherry-NC cells in midline aggregates express *mitfa*. Scale bars = 100 μm **(A)** 15 μm **(I)** and 20 μm **(N)**.

However, surprisingly, NC aggregates in *lmo7a*-deficient embryos were also positive for *foxd3* mRNA, which at this stage in WT marks Schwann cell precursors (Figure 7I-M). Furthermore, many NC cells within an aggregate clearly co-expressed both *mitfa* and *foxd3*. Wnt signaling promotes Mitf and represses Foxd3 in the context of lineage specification (Curran et al, 2010) Therefore, to assess the Wnt responses occurring in these apparent bipotential *mitfa/foxd3* double-positive NC cells we examined their expression of *7xTCF:nls-mCherry*. Although expression is mosaic in the NC aggregates of *lmo7a*-deficient embryos (Figure 7N-O), all mCherry-cells also expressed *mitfa* (Figure 7P). To investigate this further, we treated *lmo7a*-deficient embryos with a canonical Wnt inhibitor, XAV939from 12-24 hpf, during NC migration but after NC induction (Figure S3). While the treatments drastically reduced levels of the Wnt reporter (Figure S3A-F), they did not significantly alter either the number of NC cells that aggregate at the midline (DMSO mean = 28.8, XAV939 mean = 19.9, p=0.11) (Figure S3G) or that express *mitfa* (Figure S3H-M). These results suggest that elevated canonical Wnt signaling is unlikely to be the primary cause of either aberrant NC migration or the fates of NC cells in the absence of Lmo7a function and may instead be secondary to the cytoskeletal functions of Lmo7a.

## DISCUSSION

NC cells integrate many signals as they migrate to arrive at their final positions and generate the correct cell types. The gene regulatory network for NC specification and differentiation has been well studied, but the relationship between NC migration and fate is still poorly understood (Kalcheim and Kumar, 2017). Here we show novel roles for Lmo7a in regulating both NC migration and lineage specification through its functions in modulating adhesion and canonical Wnt signaling. This is the first evidence for a role for Lmo7 in NC development, and given the complex and modular nature of its function in other contexts, we can hypothesize many potential mechanisms. In our model (Figure 8), Lmo7a serves to direct formation of FAs in a similar manner to its previously described function in cultured cells (Holaska et al, 2006; Wozniak et al, 2013). Loss of Lmo7a would then interfere with formation of the Pxn-FA complex at the membrane, thereby disrupting the ability of NC cells to migrate in a coordinated manner. This is consistent with its highly restricted expression to migrating NC cells as well as its subcellular localization to the membrane where it regulates Pxn localization. We hypothesize that this loss of proper Pxn-FA formation may disrupt a complex similar to the recently described NC Cadherin11-FA complex, which includes Pxn and β-cat (Langhe et al, 2016), thereby increasing the amount of free β-cat in the cell and consequently elevating canonical Wnt signaling (Figure 8). This would be consistent with previously described roles for Lmo7 in regulating FA formation (Wozniak et al 2013) and in linking cadherins to afadin-nectin junctions (Ooshio et al, 2004).

**Figure 8.**
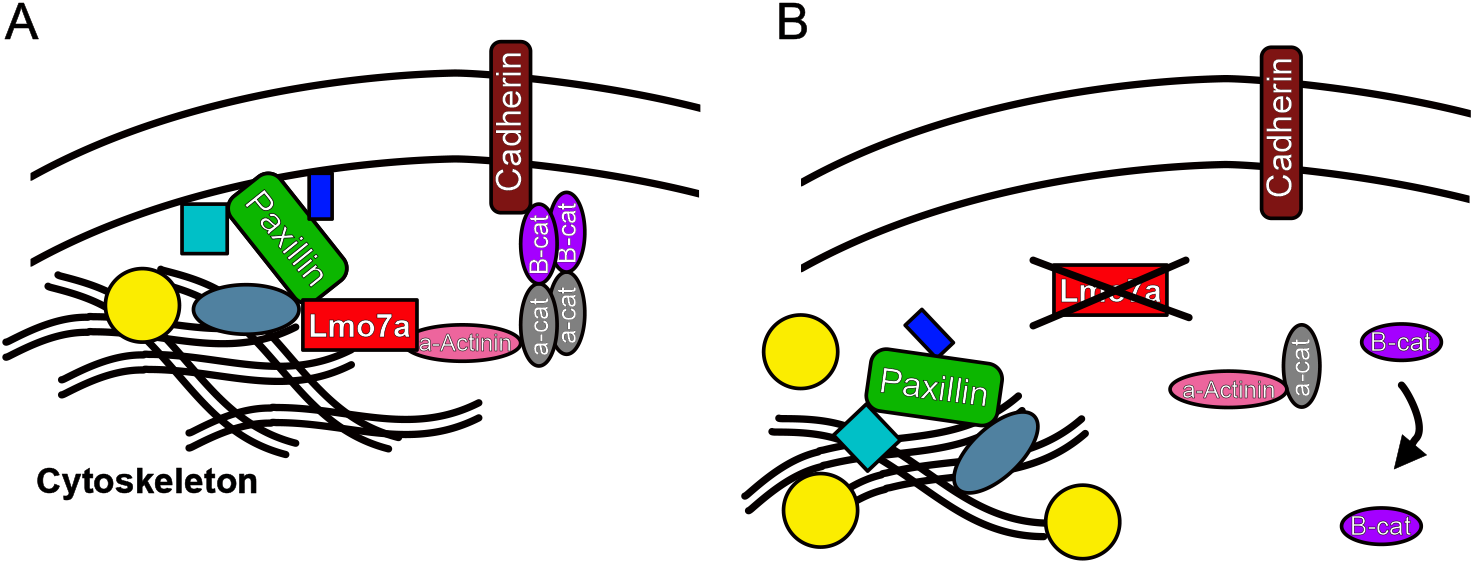
Hypothetical model for Lmo7a functions in NC cells. **(A)** Lmo7a promotes proper assembly of the Pxn-FA complex in coordination with a Cadherin-β-Cat complex similar to a classical adherens junction. Both associate with elements of the actin cytoskeleton. **(B)** In the absence of Lmo7a, the Pxn-FA complex fails to localize correctly and accumulates in the cytoplasm. This also disrupts the Cadherin-β-Cat complex, resulting in loss of directional migration and increased free β-Cat levels.

### Lmo7a regulates NC cell migration and FA localization

Lmo7 has been implicated in muscle and epithelial development as well as cancer metastasis. The mechanism of its function and its subcellular localization differ in each of these contexts. In muscle, Lmo7 interacts with transcriptional regulators in the nucleus, while in normal epithelia and breast cancer cells it interacts with transmembrane signaling molecules and the actin cytoskeleton (Holaska et al, 2006; Dedeic et al, 2011; Ooshio et al., 2004; Nakamura et al, 2005; Hu et al, 2011, Teixeira et al, 2014). A large number of alternatively spliced Lmo7 transcripts have been identified, which further increases the range of this multifaceted protein’s potential functions.

We found that an sfGFP-Lmo7a fusion protein localized to the plasma membrane of zebrafish NC cells, with no detectable expression in the nucleus. In addition,, injection of mRNA encoding an Lmo7a protein lacking the Lim domain rescued NC migration at similar levels to full length mRNA, while mRNA lacking the CH domain did not. CH domains are well known to mediate interactions with the cytoskeleton through binding with F-actin (Korenbaum and Rivero, 2002). Taken together these results strongly suggest a role for Lmo7a at the membrane, perhaps similar to its role in connecting the cytoskeleton to adherens junctions in epithelial cells (Ooshio et al, 2004).

In breast cancer cells, Lmo7 functions to modify the cytoskeleton and promote transcription of metastatic genes. Consistent with a similar structural role for Lmo7a in NC cells, we found mislocalization and aggregation of Pxn, an essential scaffolding protein for FA formation. Directional FAs have been observed in NC cells cultured in vitro (Toro-Tapia, et al, 2018), and proper FA-Integrin (Itg) signaling is crucial for directed NC migration and cardiac outflow tract formation in vivo (Dai et al, 2013). LIMCH1, a related protein with a very similar domain structure as Lmo7, controls cell migration by regulating FA formation and actomyosin dynamics (Lin et al, 2017). Lmo7 itself has been shown to localize to FAs and act as a shuttling protein to mediate Itg signaling in HeLa cells and mouse embryonic fibroblasts (Wozniak et al, 2013). The essential nature of its CH domain in NC migration supports this notion, as CH domains have been found to bind Pxn (Sjöblom et al, 2008). Our results suggest a role for Lmo7a in promoting NC cell migration through regulation of FA dynamics. This requirement appears to be specific to subsets of NC cells. This finding is similar to loss of Fscn1-dependant filopodia in premigratory NC cells, which leads to a loss of ventrolateral migration of subsets of NC cells (Boer et al, 2015). The selective loss of NC migration in *lmo7a*-deficient embryos reinforces the notion that requirements for specific migratory regulators are heterogeneous in the NC and possibly lineage-specific. This is an exciting avenue for research into the in vivo relationship between FAs and NC migration and lineage decisions.

### NC aggregates elevate Wnt signaling secondary to a loss of migratory capacity

After NC migration at 24 hpf, unmigrated NC aggregates in *lmo7a*-deficient embryos display high levels of Wnt signaling as measured using both the transgenic Wnt reporter, *Tg(7xTCF:EGFP)* and nuclear β-cat protein. This may have important consequences both for the migratory behaviors as well as the fates of the cells. By this stage most wild-type NC cells as well as migrated NC cells in *lmo7a*-deficient embryos have significantly downregulated Wnt signaling. In addition to its roles in NC induction and migration, Wnt signaling also regulates the fate decision between pigment and glial lineages. Bipotential pigment/glial progenitors downregulate *foxd3* and upregulate *mitfa* in response to elevated Wnt, biasing them toward a pigment cell fate (Curran et al, 2010). Surprisingly, despite high levels of Wnt signaling, the NC aggregates in *lmo7a*-deficient embryos express both *foxd3* and *mitfa*, often in the same cells. This is in contrast to previous findings showing that when either *rbc3a/dmxl2* or *ovol1a* are eliminated, NC cells aggregate at the dorsal midline, upregulate responses to Wnt, and all become pigment cells. Furthermore, chemical inhibition of Wnt signaling in *lmo7a*-deficient embryos fails to rescue NC migration or prevent mitfa expression in aggregates, indicating that the increase in Wnt signaling is insufficient to explain either aspect of the *lmo7a*-deficient phenotype. This is a negative result, and it is possible that the timing or effectiveness of the Wnt inhibition was not sufficient to fully block the effects of increased Wnt signaling. Comparative analysis with other known Wnt-regulatory mutants may help to disentangle the effects on migration and lineage specification.

How these functions of Lmo7a in adhesion lead to specific effects on distinct NC derivatives is less clear. Either Lmo7a functions to regulate the pigment/glial fate decision directly, or it may affect migration of a subset of NC cells that is distinct from the populations affected by *rbc3a/dmxl2* and *ovol1a*. NC aggregates in *lmo7a*-deficient embryos appear strikingly similar to loss of *rbc3a* or *ovol1*, yet distinct in that they contain both pigment and glial progenitors (Piloto and Schilling, 2009; Tuttle et al, 2014). In the case of *ovol1a*, the migration defect observed in NC cells showed a clearly correlates with elevated Wnt signaling through downstream Wnt effector genes and adoption of pigment cell fate. Similarly, in the case of *rbc3a/* dmxl2, effects on trafficking of Frizzled-7 receptors correlates with elevated nuclear β-cat and pigment cell identity. However, our results with *lmo7a* call into question a direct connection between NC aggregation, Wnt signaling and lineage specification. It is possible that this reflects either differential premigratory positioning of cells affected in each context or some yet unknown secondary transcriptional roles for one or more of these genes. Comparative transcriptomic analyses may help to resolve these differences, and further elucidate the distinct roles of these novel regulators in NC development. Our results continue to underscore the complex nature of the manifold roles for Wnt signaling in NC development.

### Conserved roles for Lmo7 in coordinating cell migration and fate

Cells that undergo EMT and migrate require a spatiotemporal balance of several interconnected cellular processes including signal transduction, actomyosin activity, and FA assembly/disassembly. Taken together, our results suggest that Lmo7a forms a novel link between these processes in migrating NC cells. While previous work has shown important roles for Lmo7 in regulating cell-cell junctions in epithelial cells (Ooshio et al, 2004; Du et al, 2019) we propose novel roles for Lmo7a in Wnt signaling in NC cells as well as in cell migration through regulation of Pxn-FAs (Figure 7). Similarly migration of mesenchymal stem cells (MSCs) and their commitment to form osteoblasts are both regulated by cytoskeletal reorganization through FAs and Itgs (Khang et al, 2012; He et al, 2016), under the control of canonical Wnt signaling (He et al, 2018).

Mammalian Lmo7 has been shown to localize to FAs and influence Itg signaling in vitro (Holaska et al, 2006; Wozniak et al, 2013) and its relative, LIMCH1, regulates FA formation and actomyosin dynamics (Lin et al, 2017). Both are associated with increased cancer metastasis (Kang et al, 2000; Furuya et al, 2002; Sasaki et al, 2003; Nakamura et al, 2005; Hu et al, 2011, Teixeira et al, 2014). While mammalian Lmo4 has been implicated in NC and cancer EMT, by directly regulating Snail and Slug, it is both structurally and functionally quite distinct from Lmo7 (Ochoa et al, 2012; Ferronha et al, 2013). Lmo7 may have gained a specific role in NC cells in regulating the association of Pxn and FAs with the actin cytoskeleton during migration, as well as linking these to Cdh and β-cat membrane/nuclear ratios, thereby indirectly influencing Wnt signaling. Future studies should examine if LMO7 plays a role in cell migration in other embryonic cells in which it is expressed (e.g. axial mesoderm that forms the notochord) and as a potential therapeutic target in cancer metastasis.

## Supporting information

Statistics Code

---

**Supplementary Movie 1. Migration of NC cells in WT embryo.** NC cells labeled with Sox10:nEOS. Embryo imaged from 12-20 hpf at 5 min intervals. NC cells migrate away from the dorsal midline and into the pharyngeal arches. Scale bar = 100 μm

**Supplementary Movie 2. Migration and formation of aggregates in *lmo7a*-deficient embryo.** NC cells labeled with Sox10:nEOS. Embryo imaged from 12-20 hpf at 5 min intervals. Most NC cells migrate away from the dorsal midline. Subsets of NC cells display non-directional movement and eventually aggregate at the midline. Scale bar = 100 μm

**Supplementary Movie 3. Paxillin accumulation in aggregate NC cells in *lmo7a*-deficient embryos.** NC cells labeled with Sox10:lyn-TdTomato and express Pxn-EGFP. Lmo7a MO-injected embryos imaged from 16-19 hpf as NC cells aggregate at the dorsal midline. Some cells display accumulation of Pxn-EGFP in the cytoplasm.

## Materials and Methods

### Zebrafish husbandry and transgenic lines

Embryos were obtained from natural breeding and staged as described in Kimmel et al, 1995. All zebrafish lines were maintained according to standard protocols (Westerfield et al, 2000). Transgenic lines used in this study include *Tg(-4.9sox10:nEOS)^w18^*(Curran et al, 2010), *Tg(-4.9sox10:lyn-tdTomato*)*^ir1040^*(Schilling et al, 2010), *Tg(-4.9sox10:lyn-EGFP)^ir866^*(Schilling et al, 2010), *Tg(−7.2sox10:EGFP)^ir937^*(Wada et al, 2005; Hoffman et al, 2010), *Tg(β-actin:Pxn-EGFP)^mai1^* (Goody et al, 2010)*, Tg(7XTCF:EGFP)^ia4^*(Moro et al, 2012), and *Tg(7XTCF:nls-mCherry)^ia5^*(Moro et al, 2012). HCR, chemical inhibition, and WNT-reporter analysis experiments were performed in double transgenic embryos derived from crosses of *Tg(7XTCF:EGFP)^ia4^* to *Tg(-4.9sox10:lyn-tdTomato)*^ir1040^. For Pxn-EGFP live imaging, *Tg(β-actin:Pxn-EGFP*)^mai1^ was crossed to *Tg(-4.9sox10:lyn-tdTomato)^ir1040^* and double-transgenic progeny were analyzed. For live imaging, embryos were mounted in 1% low-melt agarose dissolved in embryo medium containing 1% tricaine. Images were taken on either a Nikon C1 confocal or a Leica SP8 confocal. Time-lapsed movies were made on a Nikon C1 confocal using a heated stage held at 28.5°C. Images were processed using ImageJ (NIH).

### In situ hybridization and hybridization chain reaction

A 291 bp clone corresponding to a retained intron in the *lmo7a* genomic locus, was obtained from OpenBiosystems (EST BI845812) and used to generate an antisense DIG-labeled probe, which was then used for ISH as previously described (Thisse and Thisse, 2008). *foxd3*, *mitfa*, and *dlx2a* HCR probes were ordered from Molecular Technologies (Los Angeles, CA) using the accession numbers NM_131290.2, NM_130923.2, and NM_131311.2 respectively. Whole mount HCR was carried out as described (Choi et al. 2014).

### Immunohistochemistry

For β-cat staining, embryos were fixed in 4% paraformaldehyde (PFA) for 1.5 hours at room temperature, permeabilized in PBS/1%Triton-x/1%DMSO for 1 hour at room temperature (RT), washed with PBS/0.1%Triton-x/1%DMSO, and blocked with 5% Donkey Serum for 1 hour at RT. Embryos were then incubated overnight at 4°C with 1:200 β-cat antibody (GeneTex) in block. For pFAK stains, embryos embryos were fixed in 4% PFA for 1 hour at room temperature, permeabilized in acetone at −20°C for 5 minutes and then PBS/1%Triton-x/1%DMSO for 30 minutes at RT, washed with PBS/0.1%Triton-x/1%DMSO, and blocked with 5% Donkey Serum/2% BSA for 1 hour at RT. Embryos were then incubated overnight with 1:200 pFAK antibody (Invitrogen) in block. For concentrations and catalog numbers of primary and secondary antibodies see supplementary table 8.

### Molecular cloning

Coding sequences of *lmo7a* were amplified from cDNA isolated from 16 hpf zebrafish embryos. cDNA was generated using Protoscript II First Strand Synthesis Kit (New England Biolabs). Lmo7a rescue constructs and *lmo7a*:sfGFP fusion constructs were generated by Gibson Assembly (Gibson et al, 2009) into the pCS2+ vector. For primer sequences used see Supplementary Table 1.

### Morpholino and mRNA microinjections

Antisense morpholino oligonucleotide (MO) targeting *lmo7a* (5’-TCGCCACTCCATCACCGGTCAACGT-3’) and Control MO (Gene Tools) were dissolved in nanopure water prior to injection. For all mRNA injection experiments, mRNAs were transcribed in vitro using the mMessage mMachine kit (Ambion). All injections were performed at the 1-cell stage, and a volume of 1 nl was injected in all cases. For XAV939 treatment experiments, *lmo7a* MO was injected at 3 ng/embryo because of DMSO and XAV939 toxicity. For all other experiments, it was injected at 4 ng/embryo. Control MO was injected at 4 ng/embryo. In all cases, 400 pg of mRNA was injected.

### CRISPR and CRISPRi gRNA

For CRISPR-Cas9 injections, templates were generated using a different scaffold chimeric primer design (Varshney et al, 2015) and multiple target-specific primers as described (Wu et al, 2018). gRNAs were then synthesized using T7 MegaShortScript kit (Ambion). gRNAs were incubated with Cas9 protein (IDT) at 37°C for 5 minutes and injected into embryos at the 1-cell stage. 150 pg gRNA and 800 pg Cas9 protein were injected.

For CRISPRi injections, templates for gRNAs were generated by PCR using the scaffold chimeric primer design (Larson et al, 2013) and a target-specific primer containing a T7 promoter sequence. *dCas9* mRNA was synthesized from the pT3T-nls-dCas9-nls plasmid (Rossi et al, 2015) using mMessage mMachine kit (Ambion). *dCas9* mRNA was coinjected with gRNAs targeting multiple sites along the *lmo7a* coding region. 75 pg gRNAs and 450 pg *dcas9* mRNA were injected.

### Genotyping

The CRISPR-Cas9 induced Δ5 and Δ35 deletions were identified by PCR of gDNA isolated from individual juvenile fish using primers targeting Exon16 of the *lmo7a* genomic locus. Heterozygous F1 fish were incrossed to generate transheterozygous F2 fish.

### Quantitative RT-PCR

For qPCR, total RNA was extracted from biological triplicates of de-yolked CRISPRi-injected and *dCas9*-injected control embryos using trizol (Invitrogen) and purified using Direct-zol RNA Miniprep kit (Zymo). cDNA was generated using Protoscript II First Strand cDNA Synthesis kit (New England Biolabs). cDNAs were diluted 1:20 and then qPCR was carried out in triplicate using the Luna Universal qPCR Master Mix (New England Biolabs). 3 primer sets targeted different exon-exon junctions in the *lmo7a* coding region and primers targeting *rps13a* as the control housekeeping gene were used for normalization. qPCR was performed on a LightCycler 480 (Roche). Fold changes were determined by calculating ΔΔCT values for each biological replicate. For qPCR primer sequences see Supplementary Table.

### Chemical inhibition of Wnt signaling

For chemical inhibition of Wnt signaling, embryos were treated with GSK3-stabilizer XAV939 (StemCell Technologies). At 12 hpf, embryos were selected for expression of *sox10:lyn-tdTomato* and *7xTCF:EGFP* and then placed in embryo media containing either 100 μM XAV939 in 0.5mM DMSO or 0.5mM DMSO alone and incubated at 28.5°C for 12 hours.

### Statistical analyses

For mRNA rescues, medians were compared using Kruskal-Wallis ANOVA and posthoc Wilcoxon tests. For β-cat and pFAK stains, normal distribution assumptions were tested using Shapiro-Wilk tests and means were compared using Welch’s two sample T-tests. Statistical tests were carried out using R, and plots were generated in R using the ggplot2 package. Other R packages used include: plyr, dplyr, reshape2, ggpubr, ggsignif, FSA, plotly, plotrix, and gridExtra. For R markdown files including all code used for analysis and visual representation of data see Statistics Code.

## Acknowledgements

We would like to thank Michael T Green’s laboratory at UCI for providing sfGFP plasmids, Francesco Argenton’s laboratory at University of Padova for providing 7xTCF transgenics, and Clarissa Henry’s laboratory at University of Maine for providing pxn-GFP transgenics. We are grateful for the technical and critical input from members of the Schilling laboratory.

## Author Contributions

The authors have made the following declarations about their contributions:

David Tatarakis: Conceptualization, Data curation, Formal analysis, Validation, Investigation, Visualization, Methodology, Writing – original draft, Writing – review and editing, Project Administration.

Adam Tuttle: Conceptualization, Initial investigation, Writing – review and editing.

Thomas F. Schilling: Conceptualization, Resources, Data curation, Supervision, Funding acquisition, Writing – review and editing, Project Administration.

## Funding

NIH (DE023050)

NIH (DE13828)

## Supplemental Figures

**Figure S1:**
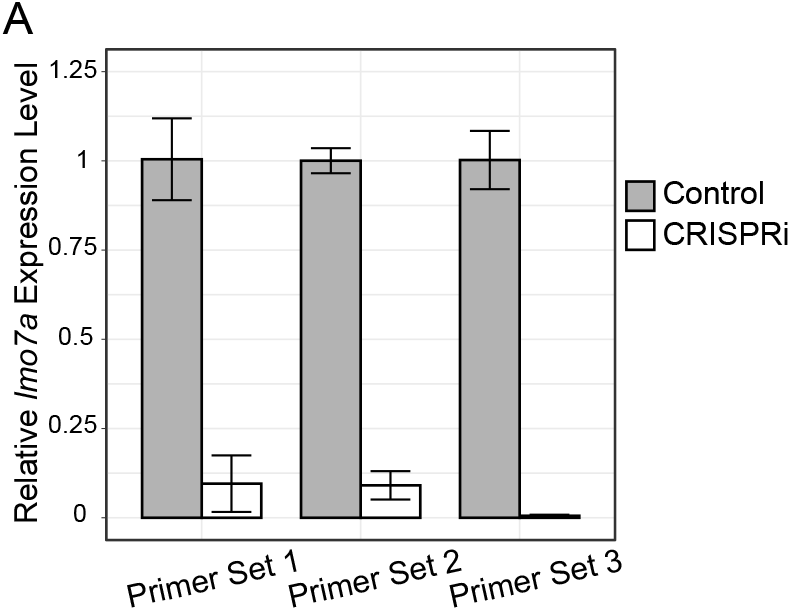
Relative expression of *lmo7a* in CRISPRi embryos. Bar plots showing expression levels of *lmo7a* based on qPCR using 3 different primer sets targeting different segments of the coding region. Bars indicate mean fold change of 3 biological replicates as compared to mean WT expression. Fold changes calculated by taking 2^ΔΔCT for each replicate. Error bars indicate mean ±SD.

**Figure S2:**
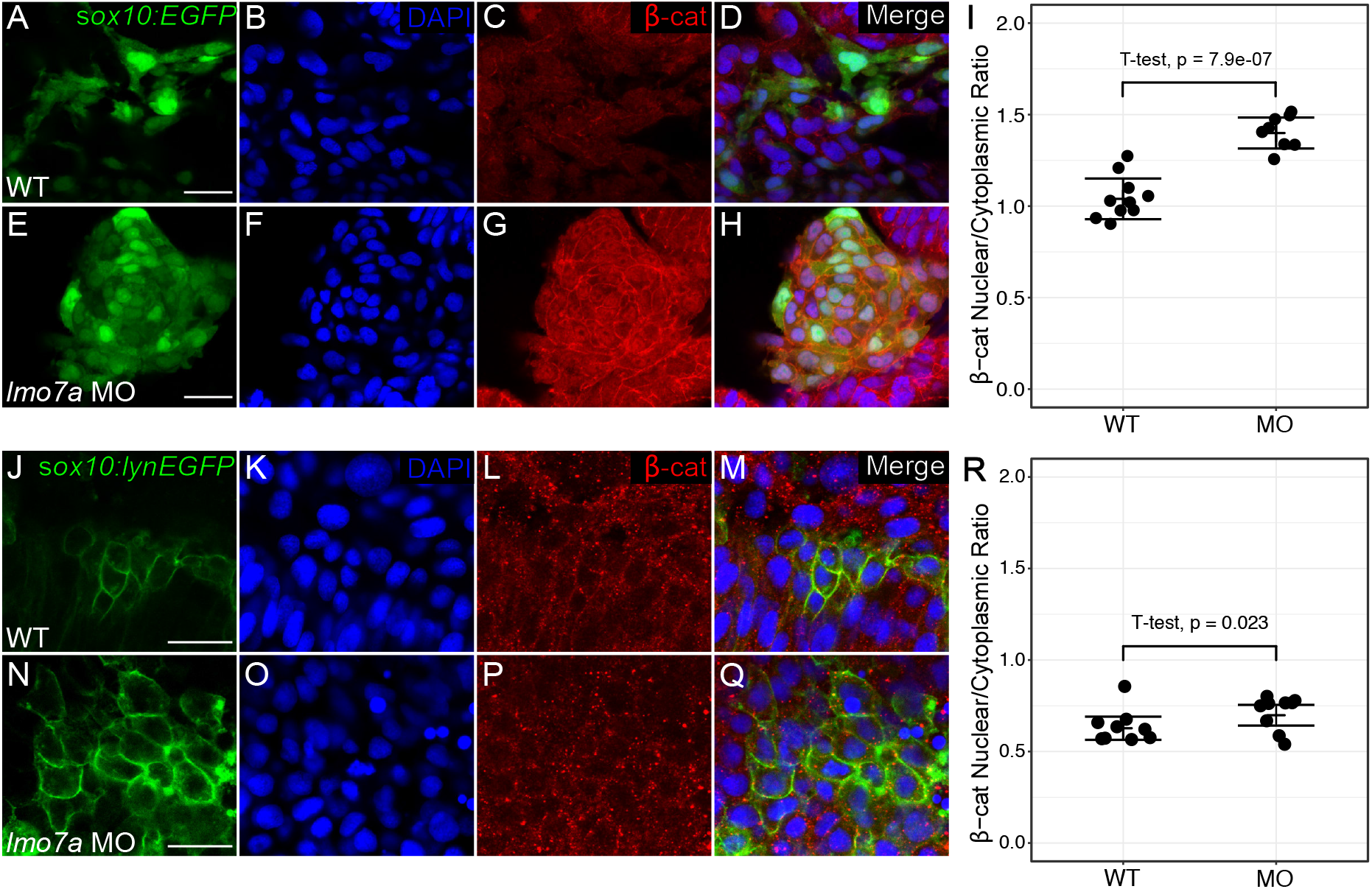
*lmo7a*-deficient NC aggregates have increased nuclear localization of β-catenin. **(A-H)** Whole mount confocal images of immunostaining with anti-β-cat antibody (red) and DAPI (blue) in 24 hpf *sox10:EGFP* transgenic embryos (green). **(A-D)** In migratory NC cells in WT embryos, β-cat staining is both in the nucleus and cytoplasm. **(E-H)** Aggregate NC cells in *lmo7a* MO-injected embryos display increased nuclear β-cat relative to cytoplasmic staining. **(I)** Nuclear β-cat levels for each cell were quantified as the mean fluorescence intensity in the nucleus divided by the mean fluorescence intensity in the cytoplasm in 10 individual cells per embryo. (WT n=10 embryos, mean=1.04; *lmo7a* MO n=8 embryos, mean=1.40). T-test p-value=7.913e-07. Line indicates mean. Error bars indicate ±SD. **(J-Q)** Whole mount confocal images of immunostaining with anti-β-cat antibody (red) and DAPI (blue) in 12 hpf *sox10:lynEGFP* transgenic embryos (green). Premigratory NC cells in WT embryos **(J-M)** and in *lmo7a* Mo-injected embryos **(N-Q)** both display low levels of β-cat staining in the nucleus. **(R)**. (WT n=9 embryos, mean=0.63; *lmo7a* MO n=9 embryos, mean=0.70). T-test p-value=0.023. Line indicates mean. Error bars indicate ±SD. Scale bars =20μm.

**Figure S3:**
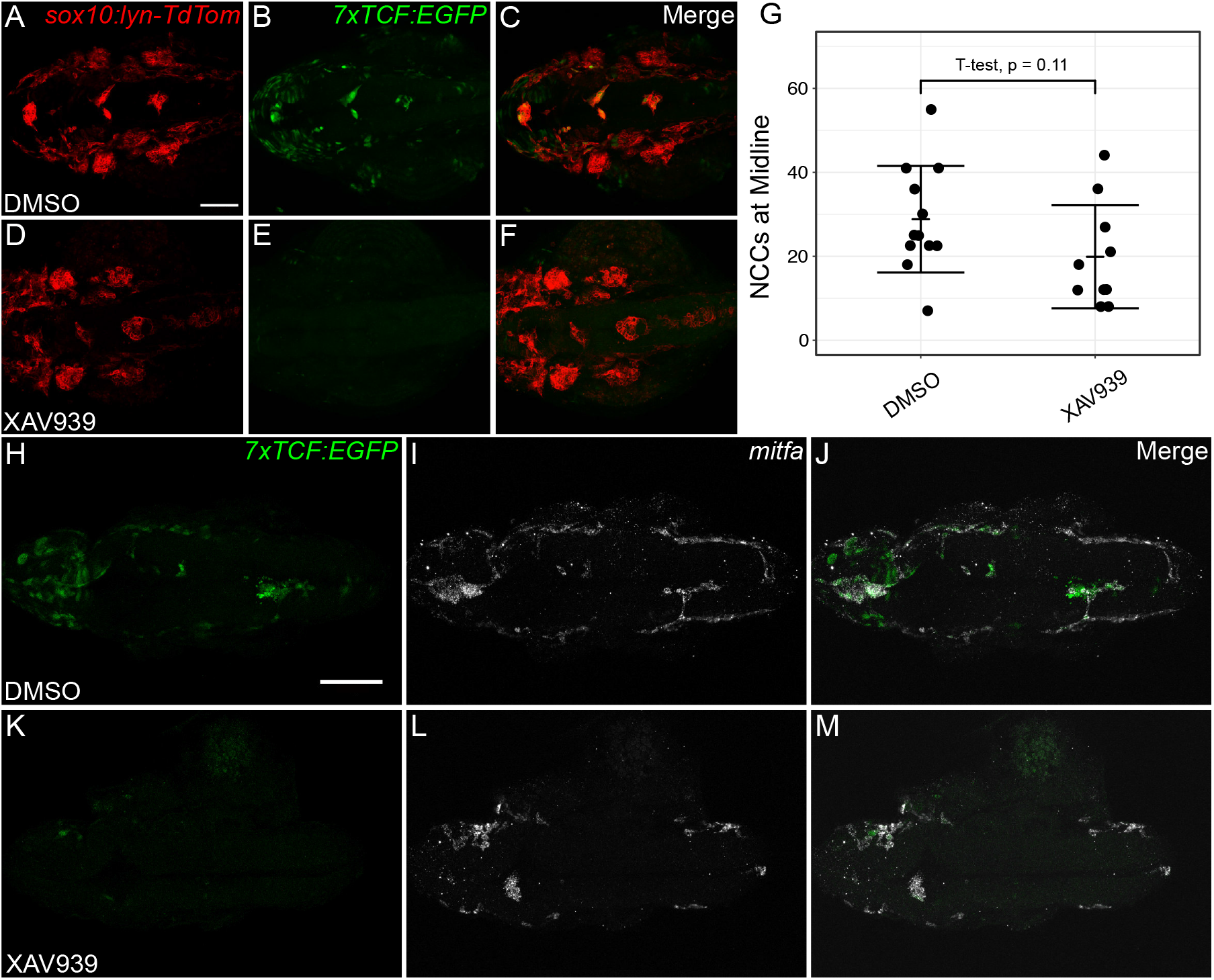
Chemical inhibition of canonical Wnt signaling does not rescue migration or block mitfa expression in *lmo7a*-deficient NC cells. **(A-F)** Whole mount live confocal images of 24 hpf *Tg(sox10:lyn-tdTomato; 7XTCF:EGFP)* double transgenic embryos. **(A-C)** In *lmo7a* MO-injected embryos treated with 0.5mM DMSO, EGFP expression is strong throughout midline NC aggregates. **(D-F)** In *lmo7a* MO-injected embryos treated with 100 μM XAV939, NC cells aggregate but lose EGFP expression. Aggregation is slightly reduced, but the change is not significant **(G)**. **(H-M)** Whole mount confocal images of HCR for *mitfa* in 24 hpf transgenic *7XTCF:EGFP* embryos. **(H-J)** In *lmo7a* MO-injected embryos treated with 0.5mM DMSO, expression of *mitfa* largely overlaps with EGFP expression. **(K-M)** In *lmo7a* MO-injected embryos treated with XAV939, *mitfa* expression persists despite nearly complete loss of EGFP expression. For **(G)** DMSO mean = 29.9 cells/embryo, XAV939 mean = 18.8 cells/embryo. T-test p-value = 0.11. Line indicates mean. Error bars indicate ±SD. Scale bars = 100 μm.

## REFERENCES

McBeath, R., Pirone, D. M., Nelson, C. M., Bhadriraju, K., & Chen, C. S. (2004). Cell Shape, Cytoskeletal Tension, and RhoA Regulate Stem Cell Lineage Commitment. Developmental Cell, 6(4), 483–495. https://doi.org/10.1016/S1534-5807(04)00075-9

He, J., Zhang, N., Zhang, J., Jiang, B., & Wu, F. (2018). Migration critically mediates osteoblastic differentiation of bone mesenchymal stem cells through activating canonical Wnt signal pathway. Colloids and Surfaces B: Biointerfaces, 171, 205–213. https://doi.org/10.1016/j.colsurfb.2018.07.017

Kalcheim, C., & Kumar, D. (2017). Cell fate decisions during neural crest ontogeny. International Journal of Developmental Biology, 61(3-4–5), 195–203. https://doi.org/10.1387/ijdb.160196ck

Stuhlmiller, T. J., & García-Castro, M. I. (2012). Current perspectives of the signaling pathways directing neural crest induction. Cellular and Molecular Life Sciences, 69(22), 3715–3737. https://doi.org/10.1007/s00018-012-0991-8

Thiery, J. P., Acloque, H., Huang, R. Y. J., & Nieto, M. A. (2009). Epithelial-Mesenchymal Transitions in Development and Disease. Cell, 139(5), 871–890. https://doi.org/10.1016/j.cell.2009.11.007

Mayor, R., & Theveneau, E. (2013). The neural crest. Development, 140(11), 2247–2251. https://doi.org/10.1242/dev.091751

Kerosuo, L., & Bronner-Fraser, M. (2012). What is bad in cancer is good in the embryo: Importance of EMT in neural crest development. Seminars in Cell & Developmental Biology, 23(3), 320–332. https://doi.org/10.1016/j.semcdb.2012.03.010

Fraser, S. E., & Bronner-Fraser, M. (1991). Migrating neural crest cells in the trunk of the avian embryo are multipotent. Development, 112(4), 913–920.

Dupin, E., Calloni, G. W., & Douarin, N. M. L. (2010). The cephalic neural crest of amniote vertebrates is composed of a large majority of precursors endowed with neural, melanocytic, chondrogenic and osteogenic potentialities. Cell Cycle, 9(2), 238–249. https://doi.org/10.4161/cc.9.2.10491

Baggiolini, A., Varum, S., Mateos, J. M., Bettosini, D., John, N., Bonalli, M., Ziegler, U., Dimou, L., Clevers, H., Furrer, R., & Sommer, L. (2015). Premigratory and migratory neural crest cells are multipotent in vivo. Cell Stem Cell, 16(3), 314–322. https://doi.org/10.1016/j.stem.2015.02.017

Schilling, T. F., & Kimmel, C. B. (1994). Segment and cell type lineage restrictions during pharyngeal arch development in the zebrafish embryo. Development, 120(3), 483–494.

Krispin, S., Nitzan, E., Kassem, Y., & Kalcheim, C. (2010). Evidence for a dynamic spatiotemporal fate map and early fate restrictions of premigratory avian neural crest. Development (Cambridge, England), 137(4), 585–595. https://doi.org/10.1242/dev.041509

Nakagawa, S., & Takeichi, M. (1998). Neural crest emigration from the neural tube depends on regulated cadherin expression. Development, 125(15), 2963–2971.

Borchers, A., David, R., & Wedlich, D. (2001). Xenopus cadherin-11 restrains cranial neural crest migration and influences neural crest specification. Development (Cambridge, England), 128(16), 3049–3060.

Morrison, J. A., McLennan, R., Wolfe, L. A., Gogol, M. M., Meier, S., McKinney, M. C., Teddy, J. M., Holmes, L., Semerad, C. L., Box, A. C., Li, H., Hall, K. E., Perera, A. G., & Kulesa, P. M. (2017). Single-cell transcriptome analysis of avian neural crest migration reveals signatures of invasion and molecular transitions. ELife, 6, e28415. https://doi.org/10.7554/eLife.28415

Lukoseviciute, M., Gavriouchkina, D., Williams, R. M., Hochgreb-Hagele, T., Senanayake, U., Chong-Morrison, V., Thongjuea, S., Repapi, E., Mead, A., & Sauka-Spengler, T. (2018). From Pioneer to Repressor: Bimodal foxd3 Activity Dynamically Remodels Neural Crest Regulatory Landscape In Vivo. Developmental Cell, 47(5), 608–628.e6. https://doi.org/10.1016/j.devcel.2018.11.009

Soldatov, R., Kaucka, M., Kastriti, M. E., Petersen, J., Chontorotzea, T., Englmaier, L., Akkuratova, N., Yang, Y., Häring, M., Dyachuk, V., Bock, C., Farlik, M., Piacentino, M. L., Boismoreau, F., Hilscher, M. M., Yokota, C., Qian, X., Nilsson, M., Bronner, M. E., … Adameyko, I. (2019). Spatiotemporal structure of cell fate decisions in murine neural crest. Science, 364(6444), eaas9536. https://doi.org/10.1126/science.aas9536

Dorsky, R. I., Moon, R. T., & Raible, D. W. (1998). Control of neural crest cell fate by the Wnt signalling pathway. Nature, 396(6709), 370–373. https://doi.org/10.1038/24620

Minchin, J. E. N., & Hughes, S. M. (2008). Sequential actions of Pax3 and Pax7 drive xanthophore development in zebrafish neural crest. Developmental Biology, 317(2), 508–522. https://doi.org/10.1016/j.ydbio.2008.02.058

Curran, K., Lister, J. A., Kunkel, G. R., Prendergast, A., Parichy, D. M., & Raible, D. W. (2010). Interplay between Foxd3 and Mitf regulates cell fate plasticity in the zebrafish neural crest. Developmental Biology, 344(1), 107–118. https://doi.org/10.1016/j.ydbio.2010.04.023

Maj, E., Künneke, L., Loresch, E., Grund, A., Melchert, J., Pieler, T., Aspelmeier, T., & Borchers, A. (2016). Controlled levels of canonical Wnt signaling are required for neural crest migration. Developmental Biology, 417(1), 77–90. https://doi.org/10.1016/j.ydbio.2016.06.022

Hutchins, E. J., & Bronner, M. E. (2018). Draxin acts as a molecular rheostat of canonical Wnt signaling to control cranial neural crest EMT. J Cell Biol, 217(10), 3683–3697. https://doi.org/10.1083/jcb.201709149

Ahsan, K., Singh, N., Rocha, M., Huang, C., & Prince, V. E. (2019). Prickle1 is required for EMT and migration of zebrafish cranial neural crest. Developmental Biology, 448(1), 16–35. https://doi.org/10.1016/j.ydbio.2019.01.018

Piloto, S., & Schilling, T. F. (2010). Ovo1 links Wnt signaling with N-cadherin localization during neural crest migration. Development, 137(12), 1981–1990. https://doi.org/10.1242/dev.048439

Tuttle, A. M., Hoffman, T. L., & Schilling, T. F. (2014). Rabconnectin-3a Regulates Vesicle Endocytosis and Canonical Wnt Signaling in Zebrafish Neural Crest Migration. PLOS Biology, 12(5), e1001852. https://doi.org/10.1371/journal.pbio.1001852

Hoffman, T. L., Javier, A. L., Campeau, S. A., Knight, R. D., & Schilling, T. F. (2007). Tfap2 transcription factors in zebrafish neural crest development and ectodermal evolution. Journal of Experimental Zoology Part B: Molecular and Developmental Evolution, 308B(5), 679–691. https://doi.org/10.1002/jez.b.21189

Li, B., Mackay, D. R., Dai, Q., Li, T. W. H., Nair, M., Fallahi, M., Schonbaum, C. P., Fantes, J., Mahowald, A. P., Waterman, M. L., Fuchs, E., & Dai, X. (2002). The LEF1/β-catenin complex activates movo1, a mouse homolog of Drosophila ovo required for epidermal appendage differentiation. Proceedings of the National Academy of Sciences, 99(9), 6064–6069. https://doi.org/10.1073/pnas.092137099

Matthews, J. M., Lester, K., Joseph, S., & Curtis, D. J. (2013). LIM-domain-only proteins in cancer. Nature Reviews Cancer, 13(2), 111–122. https://doi.org/10.1038/nrc3418

Sang, M., Ma, L., Sang, M., Zhou, X., Gao, W., & Geng, C. (2014). LIM-domain-only proteins: Multifunctional nuclear transcription coregulators that interacts with diverse proteins. Molecular Biology Reports, 41(2), 1067–1073. https://doi.org/10.1007/s11033-013-2952-1

Ochoa, S. D., Salvador, S., & LaBonne, C. (2012). The LIM adaptor protein LMO4 is an essential regulator of neural crest development. Developmental Biology, 361(2), 313–325. https://doi.org/10.1016/j.ydbio.2011.10.034

Ferronha, T., Rabadán, M. A., Gil-Guiñon, E., Dréau, G. L., Torres, C. de, & Martí, E. (2013). LMO4 is an Essential Cofactor in the Snail2-Mediated Epithelial-to-Mesenchymal Transition of Neuroblastoma and Neural Crest Cells. Journal of Neuroscience, 33(7), 2773–2783. https://doi.org/10.1523/JNEUROSCI.4511-12.2013

Martin, B., Schneider, R., Janetzky, S., Waibler, Z., Pandur, P., Kühl, M., Behrens, J., von der Mark, K., Starzinski-Powitz, A., & Wixler, V. (2002). The LIM-only protein FHL2 interacts with β-catenin and promotes differentiation of mouse myoblasts. The Journal of Cell Biology, 159(1), 113–122. https://doi.org/10.1083/jcb.200202075

Hamidouche, Z., Haÿ, E., Vaudin, P., Charbord, P., Schüle, R., Marie, P. J., & Fromigué, O. (2008). FHL2 mediates dexamethasone-induced mesenchymal cell differentiation into osteoblasts by activating Wnt/β-catenin signaling-dependent Runx2 expression. The FASEB Journal, 22(11), 3813–3822. https://doi.org/10.1096/fj.08-106302

Holaska, J. M., Rais-Bahrami, S., & Wilson, K. L. (2006). Lmo7 is an emerin-binding protein that regulates the transcription of emerin and many other muscle-relevant genes. Human Molecular Genetics, 15(23), 3459–3472. https://doi.org/10.1093/hmg/ddl423

Dedeic, Z., Cetera, M., Cohen, T. V., & Holaska, J. M. (2011). Emerin inhibits Lmo7 binding to the Pax3 and MyoD promoters and expression of myoblast proliferation genes. J Cell Sci, 124(10), 1691–1702. https://doi.org/10.1242/jcs.080259

Ooshio, T., Irie, K., Morimoto, K., Fukuhara, A., Imai, T., & Takai, Y. (2004). Involvement of LMO7 in the Association of Two Cell-Cell Adhesion Molecules, Nectin and E-cadherin, through Afadin and α-Actinin in Epithelial Cells. Journal of Biological Chemistry, 279(30), 31365–31373. https://doi.org/10.1074/jbc.M401957200

Du, T.-T., Dewey, J. B., Wagner, E. L., Cui, R., Heo, J., Park, J.-J., Francis, S. P., Perez-Reyes, E., Guillot, S. J., Sherman, N. E., Xu, W., Oghalai, J. S., Kachar, B., & Shin, J.-B. (2019). LMO7 deficiency reveals the significance of the cuticular plate for hearing function. Nature Communications, 10(1), 1–15. https://doi.org/10.1038/s41467-019-09074-4

Nakamura, H., Mukai, M., Komatsu, K., Tanaka-Okamoto, M., Itoh, Y., Ishizaki, H., Tatsuta, M., Inoue, M., & Miyoshi, J. (2005). Transforming growth factor-beta1 induces LMO7 while enhancing the invasiveness of rat ascites hepatoma cells. Cancer Letters, 220(1), 95–99. https://doi.org/10.1016/j.canlet.2004.07.023

Hu, Q., Guo, C., Li, Y., Aronow, B. J., & Zhang, J. (2011). LMO7 mediates cell-specific activation of the Rho-myocardin-related transcription factor-serum response factor pathway and plays an important role in breast cancer cell migration. Molecular and Cellular Biology, 31(16), 3223–3240. https://doi.org/10.1128/MCB.01365-10

Teixeira, V. H., Lourenco, S., Falzon, M., Capitanio, A., Bottoms, S., Carroll, B., Brown, J., George, J. P., & Janes, S. M. (2014). S112 Mmp12 And Lmo7 Are Key Genes Involved In The Early Pathogenesis Of Squamous Cell Carcinoma Of The Lung. Thorax, 69(Suppl 2), A59–A60. https://doi.org/10.1136/thoraxjnl-2014-206260.118

Lin, Y.-H., Zhen, Y.-Y., Chien, K.-Y., Lee, I.-C., Lin, W.-C., Chen, M.-Y., & Pai, L.-M. (2017). LIMCH1 regulates nonmuscle myosin-II activity and suppresses cell migration. Molecular Biology of the Cell, 28(8), 1054–1065. https://doi.org/10.1091/mbc.e15-04-0218

Wozniak, M. A., Baker, B. M., Chen, C. S., & Wilson, K. L. (2013). The emerin-binding transcription factor Lmo7 is regulated by association with p130Cas at focal adhesions. PeerJ, 1. https://doi.org/10.7717/peerj.134

Karlsson, T., Kvarnbrink, S., Holmlund, C., Botling, J., Micke, P., Henriksson, R., Johansson, M., & Hedman, H. (2018). LMO7 and LIMCH1 interact with LRIG proteins in lung cancer, with prognostic implications for early-stage disease. Lung Cancer, 125, 174–184. https://doi.org/10.1016/j.lungcan.2018.09.017

Kang, S., Xu, H., Duan, X., Liu, J. J., He, Z., Yu, F., Zhou, S., Meng, X. Q., Cao, M., & Kennedy, G. C. (2000). PCD1, a novel gene containing PDZ and LIM domains, is overexpressed in several human cancers. Cancer Research, 60(18), 5296–5302.

Furuya, M., Tsuji, N., Endoh, T., Moriai, R., Kobayashi, D., Yagihashi, A., & Watanabe, N. (2002). A novel gene containing PDZ and LIM domains, PCD1, is overexpressed in human colorectal cancer. Anticancer Research, 22(6C), 4183–4186.

Sasaki, M., Tsuji, N., Furuya, M., Kondoh, K., Kamagata, C., Kobayashi, D., Yagihashi, A., & Watanabe, N. (2003). PCD1, a novel gene containing PDZ and LIM domains, is overexpressed in human breast cancer and linked to lymph node metastasis. Anticancer Research, 23(3B), 2717–2721.

Langhe, R. P., Gudzenko, T., Bachmann, M., Becker, S. F., Gonnermann, C., Winter, C., Abbruzzese, G., Alfandari, D., Kratzer, M.-C., Franz, C. M., & Kashef, J. (2016). Cadherin-11 localizes to focal adhesions and promotes cell–substrate adhesion. Nature Communications, 7(1), 1–10. https://doi.org/10.1038/ncomms10909

Korenbaum, E., & Rivero, F. (2002). Calponin homology domains at a glance. Journal of Cell Science, 115(18), 3543–3545. https://doi.org/10.1242/jcs.00003

Toro-Tapia, G., Villaseca, S., Beyer, A., Roycroft, A., Marcellini, S., Mayor, R., & Torrejón, M. (2018). The Ric-8A/Gα13/FAK signalling cascade controls focal adhesion formation during neural crest cell migration in Xenopus. Development, 145(22). https://doi.org/10.1242/dev.164269

Sjöblom, B., Ylänne, J., & Djinović-Carugo, K. (2008). Novel structural insights into F-actin-binding and novel functions of calponin homology domains. Current Opinion in Structural Biology, 18(6), 702–708. https://doi.org/10.1016/j.sbi.2008.10.003

Dai, X., Jiang, W., Zhang, Q., Xu, L., Geng, P., Zhuang, S., Petrich, B. G., Jiang, C., Peng, L., Bhattacharya, S., Evans, S. M., Sun, Y., Chen, J., & Liang, X. (2013). Requirement for integrin-linked kinase in neural crest migration and differentiation and outflow tract morphogenesis. BMC Biology, 11, 107. https://doi.org/10.1186/1741-7007-11-107

Boer, E. F., Howell, E. D., Schilling, T. F., Jette, C. A., & Stewart, R. A. (2015). Fascin1-Dependent Filopodia are Required for Directional Migration of a Subset of Neural Crest Cells. PLOS Genetics, 11(1), e1004946. https://doi.org/10.1371/journal.pgen.1004946

Khang, D., Choi, J., Im, Y.-M., Kim, Y.-J., Jang, J.-H., Kang, S. S., Nam, T.-H., Song, J., & Park, J.-W. (2012). Role of subnano-, nano- and submicron-surface features on osteoblast differentiation of bone marrow mesenchymal stem cells. Biomaterials, 33(26), 5997–6007. https://doi.org/10.1016/j.biomaterials.2012.05.005

Westerfield (2000). The zebrafish book. A guide for the laboratory use of zebrafish (Danio rerio). Eugene: University of Oregon Press.

Goody, M. F., Kelly, M. W., Lessard, K. N., Khalil, A., & Henry, C. A. (2010). Nrk2b-mediated NAD+ production regulates cell adhesion and is required for muscle morphogenesis in vivo: Nrk2b and NAD+ in muscle morphogenesis. Developmental Biology, 344(2), 809–826. https://doi.org/10.1016/j.ydbio.2010.05.513

Wada, N., Javidan, Y., Nelson, S., Carney, T. J., Kelsh, R. N., & Schilling, T. F. (2005). Hedgehog signaling is required for cranial neural crest morphogenesis and chondrogenesis at the midline in the zebrafish skull. Development, 132(17), 3977–3988. https://doi.org/10.1242/dev.01943

Schilling, T. F., Pabic, P. L., & Hoffman, T. L. (2010). Using transgenic zebrafish (Danio rerio) to study development of the craniofacial skeleton. Journal of Applied Ichthyology, 26(2), 183–186. https://doi.org/10.1111/j.1439-0426.2010.01401.x

Moro, E., Ozhan-Kizil, G., Mongera, A., Beis, D., Wierzbicki, C., Young, R. M., Bournele, D., Domenichini, A., Valdivia, L. E., Lum, L., Chen, C., Amatruda, J. F., Tiso, N., Weidinger, G., & Argenton, F. (2012). In vivo Wnt signaling tracing through a transgenic biosensor fish reveals novel activity domains. Developmental Biology, 366(2), 327–340. https://doi.org/10.1016/j.ydbio.2012.03.023

Choi, H. M. T., Beck, V. A., & Pierce, N. A. (2014). Next-Generation in Situ Hybridization Chain Reaction: Higher Gain, Lower Cost, Greater Durability. ACS Nano, 8(5), 4284–4294. https://doi.org/10.1021/nn405717p

Gibson, D. G., Young, L., Chuang, R.-Y., Venter, J. C., Hutchison, C. A., & Smith, H. O. (2009). Enzymatic assembly of DNA molecules up to several hundred kilobases. Nature Methods, 6(5), 343–345. https://doi.org/10.1038/nmeth.1318

Larson, M. H., Gilbert, L. A., Wang, X., Lim, W. A., Weissman, J. S., & Qi, L. S. (2013). CRISPR interference (CRISPRi) for sequence-specific control of gene expression. Nature Protocols, 8(11), 2180–2196. https://doi.org/10.1038/nprot.2013.132

Varshney, G. K., Pei, W., LaFave, M. C., Idol, J., Xu, L., Gallardo, V., Carrington, B., Bishop, K., Jones, M., Li, M., Harper, U., Huang, S. C., Prakash, A., Chen, W., Sood, R., Ledin, J., & Burgess, S. M. (2015). High-throughput gene targeting and phenotyping in zebrafish using CRISPR/Cas9. Genome Research, 25(7), 1030–1042. https://doi.org/10.1101/gr.186379.114

Wu, R. S., Lam, I. I., Clay, H., Duong, D. N., Deo, R. C., & Coughlin, S. R. (2018). A Rapid Method for Directed Gene Knockout for Screening in G0 Zebrafish. Developmental Cell, 46(1), 112–125.e4. https://doi.org/10.1016/j.devcel.2018.06.003

Rossi, A., Kontarakis, Z., Gerri, C., Nolte, H., Hölper, S., Krüger, M., & Stainier, D. Y. R. (2015). Genetic compensation induced by deleterious mutations but not gene knockdowns. Nature, 524(7564), 230–233. https://doi.org/10.1038/nature14580

